# Chromosome-level genome reference of *Venturia effusa*, causative agent of pecan scab

**DOI:** 10.1101/746198

**Authors:** David J. Winter, Nikki D. Charlton, Nick Krom, Jason Shiller, Clive H. Bock, Murray P. Cox, Carolyn A. Young

**Affiliations:** School of Fundamental Sciences and the Bio-Protection Research Centre, Massey University, Palmerston North 4442, New Zealand; Noble Research Institute, LLC, Ardmore, OK 73401; United States Department of Agriculture – Agricultural Research Service – Southeastern Fruit and Tree Nut Research Laboratory, Byron, GA 31008

## Abstract

Pecan scab, caused by *Venturia effusa*, is a devastating disease of pecan (*Carya illinoinensis*), which results in economic losses on susceptible cultivars throughout the southeastern U.S. To enhance our understanding of pathogenicity in *V. effusa*, we have generated a complete telomere-to-telomere genome reference of *V. effusa* isolate FRT5LL7-Albino. By combining Illumina MiSeq and Oxford Nanopore MinION data, we assembled a 45.2 Mb genome represented by 20 chromosomes and containing 10,820 genes, of which 7,619 have at least one functional annotation. The likely causative mutation of the albino phenotype was identified as a single base insertion and a resulting frameshift in the gene encoding the polyketide synthase *ALM1*. This genome represents the first full chromosome level assembly of any *Venturia* species.

## Genome announcement

*Venturia* are widespread fungal pathogens that cause scab disease on pecan (*V. effusa*), apples (*V. inaequalis*), pear (*V. pyrina* and *V. nashicola*) and peach (*V. carpophila*) (González-Domínguez et al. 2017). Pecan scab is polycyclic during the growing season and most prominent on leaves, twigs and shucks. Pecan scab-susceptible cultivars are managed by fungicide application throughout the growing season to reduce economic losses due to reduced nut size and number, and poor nutmeat quality as a result of the disease (Bock et al. 2017a). *V. effusa* was previously assumed to reproduce solely asexually, but the recent identification of a heterothallic mating system and the fact that the mating types are in equilibrium throughout orchards in the southeastern USA, indicates a sexual stage may contribute to the disease cycle (Bock et al. 2018; Young et al. 2018). At present, there are more than 90 genome assemblies available for *Venturia* species, representing an active research area of fungal genomics. However, many of the genomes are highly fragmented ranging from 66 scaffolds to more than 40,000 (Bock et al. 2016b; Chen et al. 2017; Deng et al. 2017; Passey et al. 2018; Johnson et al. 2019; Le Cam et al. 2019; Prokchorchik et al. 2019). Here, we present the first entire and finished genome sequence for the genus *Venturia*.

Isolate FRT5LL7 was originally collected on 23 June 2015 from a commercial orchard of the pecan cultivar ‘Desirable’ near southwest Albany, GA at the Graham Pecan Farm. The pecan orchard received typical pecan management for Georgia (Wells 2019). FRT5LL7 was isolated from a lesion on a leaflet in the lower tree canopy (5-8 m). FRT5LL7 was a regular melanized scab isolate that spontaneously produced an albino sector once cultured on potato dextrose agar. The albino form of FRT5LL7 (FRT5LL7-Albino) was isolated on 4 September 2015 and maintained as a separate culture. FRT5LL7-Albino was chosen for genome sequencing as the quality of the extracted DNA was superior to that of melanized isolates. In addition, FRT5LL7-Albino has been used in sexual crosses, where it has provided a useful phenotype for determining recombination frequencies during sexual crosses (Young, Charlton and Bock, unpublished). No other phenotypic anomalies of FRT5LL7-Albino have been noted.

The FRT5LL7-Albino genome was sequenced using both short- and long-read chemistry with Illumina MiSeq and Oxford Nanopore MinION, respectively. Genomic DNA was extracted from lyophilized mycelium using the Quick Fungal/Bacterial miniprep kit (Zymo Research) (omitting the mycelial pulverization with the beads) for Illumina MiSeq sequencing by the Bluegrass Technical and Community College (Lexington, KY). The Taha DNA extraction method (Al-Samarrai and Schmid 2000) was used on the same mycelium for Oxford Nanopore MinION sequencing. DNA samples were purified using a SPRI bead mixture (Schalamun and Schwessinger 2017) to reduce the number of short DNA fragments. The library for MinION sequencing was prepared using the 1D library SQK-LSK09 kit and sequenced on one R9.4 MinION flowcell with the basecalling option turned on.

Initial assemblies were produced from the basecalled nanopore data using two programs with alternative assembly algorithms, CANU v 1.8 (Koren et al. 2017) and NECAT v 20190307 (Xiao et al. 2017). The different algorithms complimented each other, with discontinuities present in one assembly being well-resolved in the other. The final assembly was polished with Pilon v 1.23 (Walker et al. 2014), using 250 bp PE Illumina MiSeq reads as input. The assembly of FRT5LL7-Albino contains 20 complete chromosomes totaling 45.2 Mb of sequence and a circular mitochondrial genome (139 kb). This is a finished genome reference; it stretches from telomere-to-telomere and contains no ambiguity bases or gaps. The 20 chromosomes range in size from the largest at 4.12 Mb to the smallest at 0.56 Mb (Figure 1). The FRT5LL7-Albino assembled genome is a vast improvement over the earlier sequenced genome of *V. effusa* isolate 3Des10b, which consists of 40.6 Mb represented by 545 scaffolds (Bock et al. 2016b).

**Figure 1.**
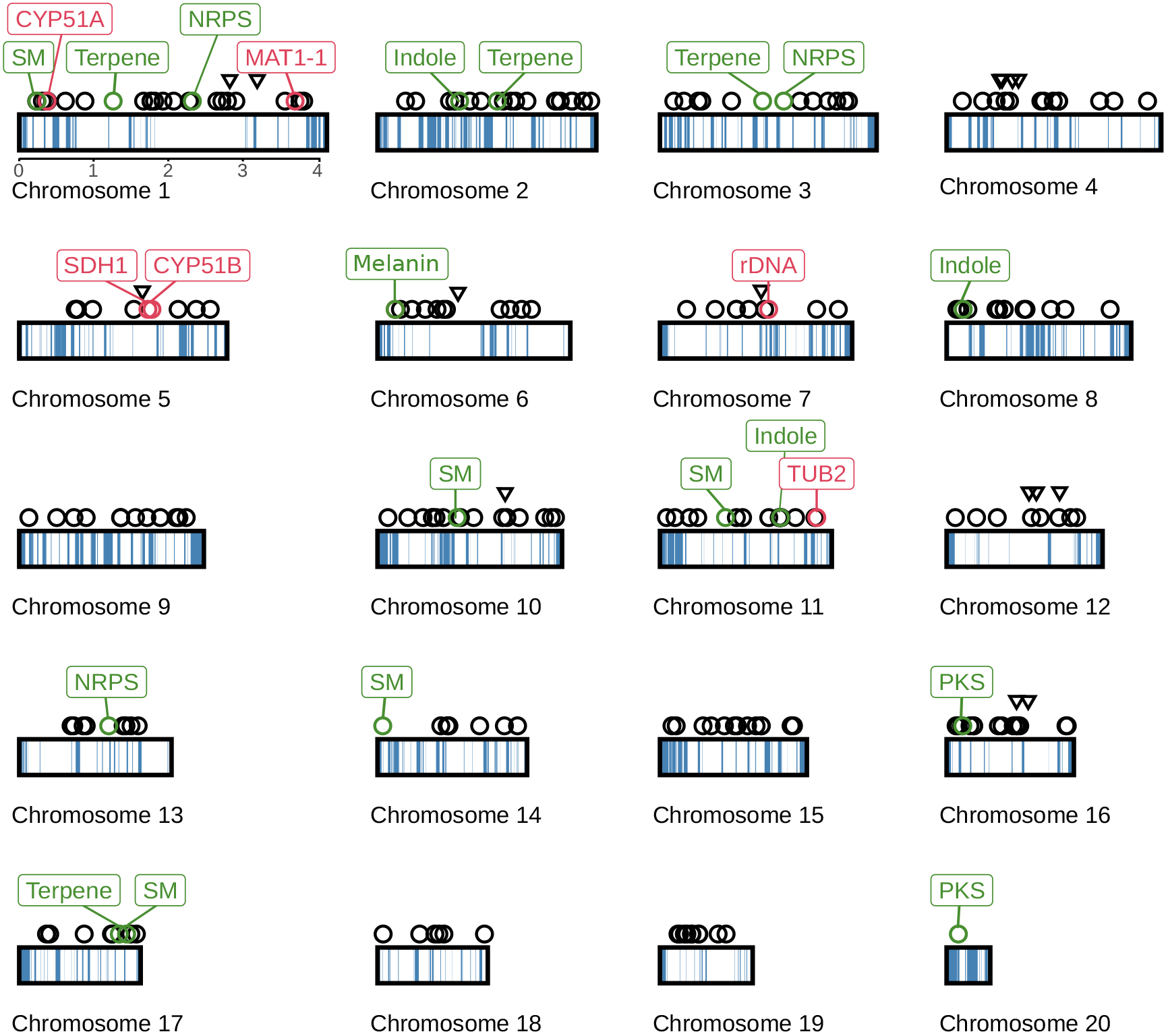
Features of the complete *Venturia effusa* genome. Each chromosome is plotted as a rectangle with repeat-dominated AT-rich regions shaded in blue. Above, key loci are indicated including secondary metabolite clusters (green circles), putative effectors (black circles), SSR loci (Bock et al. 2016a; Bock et al. 2017b) (black triangles) and other widely-studied genes such as mating type (red circles). Secondary metabolite clusters are labeled according to the key genes included in each cluster: ‘Terpene’ = terpene synthase, ‘Indole’ = aromatic prenyltransferase (e.g. dimethylallyltryptophan synthase), ‘NRPS’ = non-ribosomal peptide synthetase, ‘PKS’ = polyketide synthase, and ‘melanin’ = a cluster of three genes, *ALM1* (encoding a PKS), *CRM1* (encoding a C2H2 transcription factor) and *BRM2* (encoding a THN reductase) that encode steps involved in melanin biosynthesis.

The genome was annotated using the funannotate pipeline (v 1.4) (Palmer), which makes use of AUGUSTUS v 3.3.1 (Stanke et al. 2008) for gene-calling. Having identified putative genes, Funannotate uses InterProScan v 5.22 (Jones et al. 2014), antismash v 4.20 (Blin et al. 2017), MEROPS (Rawlings et al. 2007), Pfam (Bateman et al. 2004) and dbCAN (Huang et al. 2017) databases to generate functional annotation. In addition, we identified putative effector genes using SignalP v 4.1 (Petersen et al. 2011) and EffectorP v 2.0 (Sperschneider et al. 2016), and located AT-rich regions of the genome using OcculterCut v 1.1 (Testa et al. 2016). Our annotated genome contains 10,820 protein coding genes, of which 7,619 have at least one functional annotation. This includes 4,749 genes associated with Gene Ontology terms, 426 putative effectors and 207 genes associated with secondary metaboliite clusters (Supplementary Table S1). The BUSCO score was 96.8%, with 1272 complete BUSCOs from 1315 identified. The full functional annotation of the genome is included in Supplementary Table S2.

The higher-level structure of the genome is characterized by large blocks of AT-rich sequences interspersed between regions of approximately equal nucleotide content. The AT-rich regions are extremely gene-poor, containing only 0.15% of the protein coding genes despite comprising 24.7% of the genome. This is similar to the genome structure seen in other *Venturia* species, most notably the genomes of *V. inaequalis*, which on average have 45.3% AT-rich sequence (Le Cam et al. 2019). This type of genome structure was first described in fungi for *Leptosphaeria maculans* (Rouxel et al. 2011). Subsequently, the evolution of effector genes underlying gene-for-gene interactions in the *L. maculans-Brassica Napus* pathosystem has been shown to be associated with their proximity to or position within AT-rich regions (Van de Wouw et al. 2010; Rouxel et al. 2011). Gene-for-gene interactions have been described for other *Venturia* species (Bus et al. 2011; Deng et al. 2017) and effectors are likely to be important for the *V. effusa*-*C. illinoinensis* interaction. A gapless genome assembly of *V. effusa* spanning complete AT-rich regions is crucial to identify and understand the evolution of effector genes in the species. Key features of the genome, including AT-rich regions and the location of important genes, are illustrated in Figure 1.

The albino phenotype is often due to mutation of a melanin biosynthesis gene. Evaluation of the putative melanin biosynthesis gene cluster found on chromosome 6 in FRT5LL7-Albino (Figure 1) revealed a single base insertion and a resulting frameshift in the gene encoding the polyketide synthetase *ALM1*. As found with other melanin producing fungi, *ALM1* in *V. effusa* is clustered with two genes, *CRM1* and *BRM2* that encode putative functions in melanin biosynthesis, with an additional gene, *SDH1*, located on chromosome 5 (Kimura and Tsuge 1993; Fetzner et al. 2014).

Although other *Venturia* genomes have been sequenced (Bock et al. 2016b; Chen et al. 2017; Deng et al. 2017; Passey et al. 2018; Johnson et al. 2019; Le Cam et al. 2019; Prokchorchik et al. 2019), the complete genome sequence of *V. effusa* FRT5LL7-Albino represents the first chromosomal level assembly of a *Venturia* species. This valuable resource is a basis for scientists in the *Venturia* research community to advance the comparative genomics of scab pathogens; it is a useful resource for the identification of effectors, will provide insights into the epidemiology and population biology of the pathogens, and the evolution of the genus *Venturia*.

The genome sequence has been deposited into GenBank under genome accession CP042185 (BioProject PRJNA551043, BioSample SAMN12143997, and SRA Accessions SRX6745934 and SRX6745935 for Illumina and Nanopore MinION data respectively).

## Acknowledgments

We would like to thank Megan McDonald (Australian National University) and Simona Florea (University of Kentucky) for their excellent advice and guidance before embarking on the MinION sequencing.

This article reports the results of research only. Mention of a trademark or proprietary product is solely for the purpose of providing specific information and does not constitute a guarantee or warranty of the product by the U.S. Department of Agriculture and does not imply its approval to the exclusion of other products that may also be suitable.

---

**Supplementary Table S1.** Gene annotation statistics. “Annotation” refers to the category of functional annotation provided by the funannotate pipeline. “SMCOG” refers to genes predicted to have secondary metabolite activities, while “antiSMASH” includes all genes that form part of a putative secondary metabolite cluster. “Name” and “product” signify genes that have been assigned an NCBI gene name or protein product based on homology to well-characterized genes. The “genes annotated” column provides the number of genes having at least one annotation of each type in our dataset.

| **Annotation** | **Genes annotated** |
| --- | --- |
| Gene | 10822 |
| InterPro | 7619 |
| PFAM | 6777 |
| GO function | 4749 |
| BUSCO | 1273 |
| signal peptide | 1130 |
| Product | 583 |
| Name | 564 |
| EffectorP | 426 |
| CAZy | 405 |
| MEROPS | 320 |
| antiSMASH | 207 |
| SMCOG | 49 |

Supplementary Table S2

Functional annotations assigned to each gene. The “gene” column provides the GenBank identifier for a given gene. All other columns provide functional annotations associated with this gene. In most cases, the value provided in an annotation column is the unique identifier for protein domain, functional ontology or protein-type. When a single gene has more than one annotation of single type the unique IDs are separated by colons. The “secreted” column provides the location of signal a predicted signal peptide in a given protein product, “effectorP” identifies small secreted proteins that are putative effectors, and “SM Cluster” assigns genes to one of 18 secondary metabolite clusters.

